# Testing diffusion-derived orientation priors for streamline modelling of the MRI-visible glioblastoma core: the BRIAN framework

**DOI:** 10.64898/2026.07.30.741684

**Authors:** Aitor Alberdi Escudero, Kenneth Scerri, Andrew Sammut, Claude J. Bajada

## Abstract

Glioblastoma spreads diffusely beyond the abnormality visible on conventional magnetic resonance imaging, and because tumour cells migrate preferentially along white-matter pathways, growth models that assume isotropic spread may misrepresent the geometry of invasion relevant to radiotherapy planning. This work introduces and evaluates BRIAN, a diffusion-informed simulator that biases tumour propagation along directions derived from diffusion MRI. Patient tumour masks from the UPENN-GBM cohort are transferred onto healthy host brains from the Human Connectome Project through the MNI152 template as a proxy, diffusion orientation distribution functions (dODFs) are reconstructed on each host, and stochastic streamline propagation with the MRtrix3 iFOD2 algorithm yields volumetric occupancy maps. Propagation parameters are fitted per tumour under a three-stage curriculum that tightens a constrained five-metric objective. Across 30 unifocal tumours, each propagated onto 65 validation hosts (1 950 tumour–host pairings), the simulator reached a per-tumour median Dice coefficient of 0.748 (0.745 pooled across all pairings) and a median bounded Hausdorff agreement of 0.776. Because parameters are calibrated against each tumour’s own reference mask and the train/validation split is over hosts, these figures quantify reproduction and host-transfer of a known lesion; prediction of unseen tumours is outside their scope. On a purposively selected ten-tumour subset, controlled comparisons tested both the contribution of directional information and whether a richer angular reconstruction improved performance. Relative to a direction-blind isotropic null, orientation-informed tracking improved all five evaluated metrics (*d_z_* = 0.4–1.2), although only surface Dice remained significant after Holm correction. A second comparison replaced the multi-shell SHORE dODF with a single-tensor dODF while keeping the tracking framework unchanged. No statistically detectable differences were observed between the two directional models on this subset, providing no evidence that resolving crossing fibres improved agreement with the visible tumour envelope. Directional sampling improved agreement with the MRI-visible tumour core, but the selected subset provided no evidence that the SHORE dODF outperformed the tensor dODF.

## 1 Introduction

Glioblastoma (GBM) is the most common malignant primary brain tumour in adults and one of the most lethal human cancers, and outcomes remain poor under standard management with resection, radiotherapy, and temozolomide (Wen et al., 2020; Stupp et al., 2005; Weller et al., 2021). The dominant reason is biological: glioblastoma infiltrates diffusely, so tumour cells extend well beyond the abnormality visible on conventional MRI, and this occult disease drives local recurrence (Claes et al., 2007). Radiotherapy responds by expanding a clinical target volume (CTV) around the visible lesion to cover microscopic spread, but margin choices vary across protocols, and larger margins expose a greater volume of normal brain to radiation, increasing the risk of treatment-related injury (Niyazi et al., 2016, 2023; Lawrence et al., 2010). Therefore, the operational question that patient-specific growth modelling tries to answer is where tumour cells are likely to extend beyond the radiologically visible margin.

Glioma invasion is not spatially uniform. Tumour cells exploit existing anatomical routes, including white-matter pathways that offer low-resistance trajectories, producing anisotropic spread that isotropic growth models cannot reproduce (Claes et al., 2007; Cuddapah et al., 2014). Diffusion MRI (dMRI) is well suited to capturing this structure, because it infers tissue microstructural organisation non-invasively from the orientation dependence of water diffusion, and peritumoural diffusion abnormalities consistent with tract involvement suggest it captures features of preferential invasion routes that conventional imaging misses (Alexander et al., 2007; Price et al., 2003).

The classical proliferation–invasion framework models tumour-cell proliferation and tissue-dependent motility using scalar diffusion coefficients (Swanson et al., 2000; Harpold et al., 2007). Later models introduced anisotropic operators derived from diffusion tensors (Jbabdi et al., 2005; Clatz et al., 2005; Painter and Hillen, 2013). A structural feature runs through this literature, including related geometric and multiscale formulations (Konukoglu et al., 2010; Engwer et al., 2015): the directional input is a single second-order tensor, which encodes only one dominant orientation per voxel and so cannot represent crossing, kissing, branching, or fanning fibre populations (Tuch, 2004). Collapsing the local angular profile to one axis discards candidate directions that a directional prior on invasion would otherwise retain.

Richer angular reconstructions were developed to recover this structure. High-angular-resolution diffusion imaging supports the dODF, which resolves the multiple intravoxel orientations a single tensor collapses to one axis (Tuch, 2004; Aganj et al., 2010). Existing diffusion-informed glioma models commonly use a single tensor direction, despite diffusion MRI supporting multiple intravoxel orientations that can be followed probabilistically.

BRIAN uses diffusion-derived orientation distributions to guide stochastic streamline propagation from within the tumour. Streamlines are generated using the iFOD2 algorithm implemented in MRtrix3 and converted into volumetric occupancy maps (Tournier et al., 2010, 2019). The simulator is calibrated per tumour against the observed mask through a multi-metric, curriculum-driven parameter search whose metrics are organised under the *Metrics Reloaded* taxonomy (Maier-Hein et al., 2024). Because diffusion MRI acquired before tumour development was unavailable for the patients, patient tumour masks were transferred through MNI152 space onto healthy HCP brains (Van Essen et al., 2013; Glasser et al., 2013). The diffusion field of each healthy host was used as a proxy for the patient’s pre-tumour tissue organisation.

BRIAN was evaluated on 30 glioblastomas from the UPENN-GBM cohort (Bakas et al., 2022). Each tumour was simulated on 20 training and 65 validation host brains, with the split defined over hosts. The main comparison tested whether directional sampling improved agreement with the MRI-visible tumour core by contrasting the SHORE dODF condition with an isotropic null. A secondary comparison tested whether the multi-shell SHORE dODF improved performance over a single-tensor dODF under the same tracking framework.

## 2 Methods

The inputs were pre-operative tumour segmentations from UPENN-GBM and T1-weighted and multi-shell diffusion MRI from HCP. For each tumour–host pair, the patient mask was transferred to the host brain through MNI152 space, streamlines were propagated using the host diffusion field, and the resulting occupancy map was converted into a binary simulated mask. Propagation parameters were fitted to the reference tumour using a multi-metric optimisation procedure.

### 2.1 Data and case selection

Healthy host brains were drawn from the *100 Unrelated Subjects* package of the HCP 1200 Subjects release (Van Essen et al., 2013; Glasser et al., 2013), taken as distributed, corresponding to the HCP minimally preprocessed pipelines (Glasser et al., 2013). The structural channel is a 3D T1-weighted MP-RAGE at 0.7 mm isotropic; the diffusion channel is multi-shell at *b* = 1000, 2000 and 3000 s/mm^2^ with 90 directions per shell at 1.25 mm isotropic (Van Essen et al., 2013; Sotiropoulos et al., 2013; Glasser et al., 2013). The T1-weighted volume, the diffusion data and gradient tables, and the native-space brain mask were used. Quality control used the distributed quality control issues table, generated as part of the HCP quality-control infrastructure (Marcus et al., 2013). For the present study, subjects with a recorded QC issue were excluded, leaving 86 subjects; one further subject whose local data could not be read was excluded, leaving 85 usable hosts. Of these, 20 were assigned to optimiser training and 65 to validation, the same split for every tumour. Because the split is over disjoint host pools, every tumour appears in both roles while no validation host contributes to the parameters used to simulate it.

Patient tumour masks came from the University of Pennsylvania Glioblastoma (UPENNGBM) dataset (Bakas et al., 2022), which provides pre-operative multi-parametric MRI with automated segmentations refined by expert review, together with per-subject molecular annotations that include IDH1 status. Structural scans are distributed after the Brain Tumor Segmentation (BraTS) preprocessing pipeline, that is, with only the standardised minimal preprocessing that pipeline applies: rigid co-registration to the SRI24 atlas, skull-stripping, and resampling to 1 mm isotropic (Bakas et al., 2022). The native T1-weighted image (the moving image in the UPENN-to-template registration) and the automated segmentation were used. The target object was the tumour core, labels 1 and 4 in the native segmentation (Bakas et al., 2022), combined by voxel-wise maximum and binarised into a single foreground mask; peritumoural oedema was excluded because the reference mask represents the tumour core. A set of 30 tumours was selected by sequential visual inspection of the binarised masks in cohort-identifier order, retaining only cases whose mask formed a single connected component. Cases where the union of labels 1 and 4 produced two or more disconnected components (multifocal disease or detached satellites) were excluded, since a single seeded propagation cannot cover separated foci and would inflate false positives between them. The resulting cohort spans a range of volumes, hemispheric and lobar locations, and morphologies. Selection did not filter on molecular status; of the 30 tumours, 28 are recorded as IDH-wildtype and two as IDH-NOS/NEC (status not determined) in the UPENN-GBM clinical metadata (Bakas et al., 2022). The study uses de-identified, publicly released data; ethics approval was obtained from the Faculty of Medicine and Surgery Research Ethics Committee of the University of Malta (MED-2025-00309).

### 2.2 Registration

No diffusion MRI exists for the pre-diagnostic healthy state of a patient’s own brain, so the method decouples tumour shape, taken from the patient, from the microstructural environment, taken from a healthy HCP subject; this requires transferring the patient tumour mask from native UPENN T1 space into native HCP T1 space. In preliminary testing, registering the healthy HCP T1 directly onto the patient UPENN T1 and inverting the result could place portions of tumours outside the host brain, particularly near cortical or ventricular boundaries. An intermediate reference was therefore used, the MNI152 NLin2009cAsym template at 1 mm isotropic in brain-extracted form (Fonov et al., 2009, 2011), a standard common space for inter- subject analysis (Evans et al., 2012). Patient and host are registered into MNI independently, and the composition of the two registrations, one inverted, carries the tumour from UPENN T1 to HCP T1 without forcing the host brain to match the tumour region (Figure 1).

**Figure 1:**
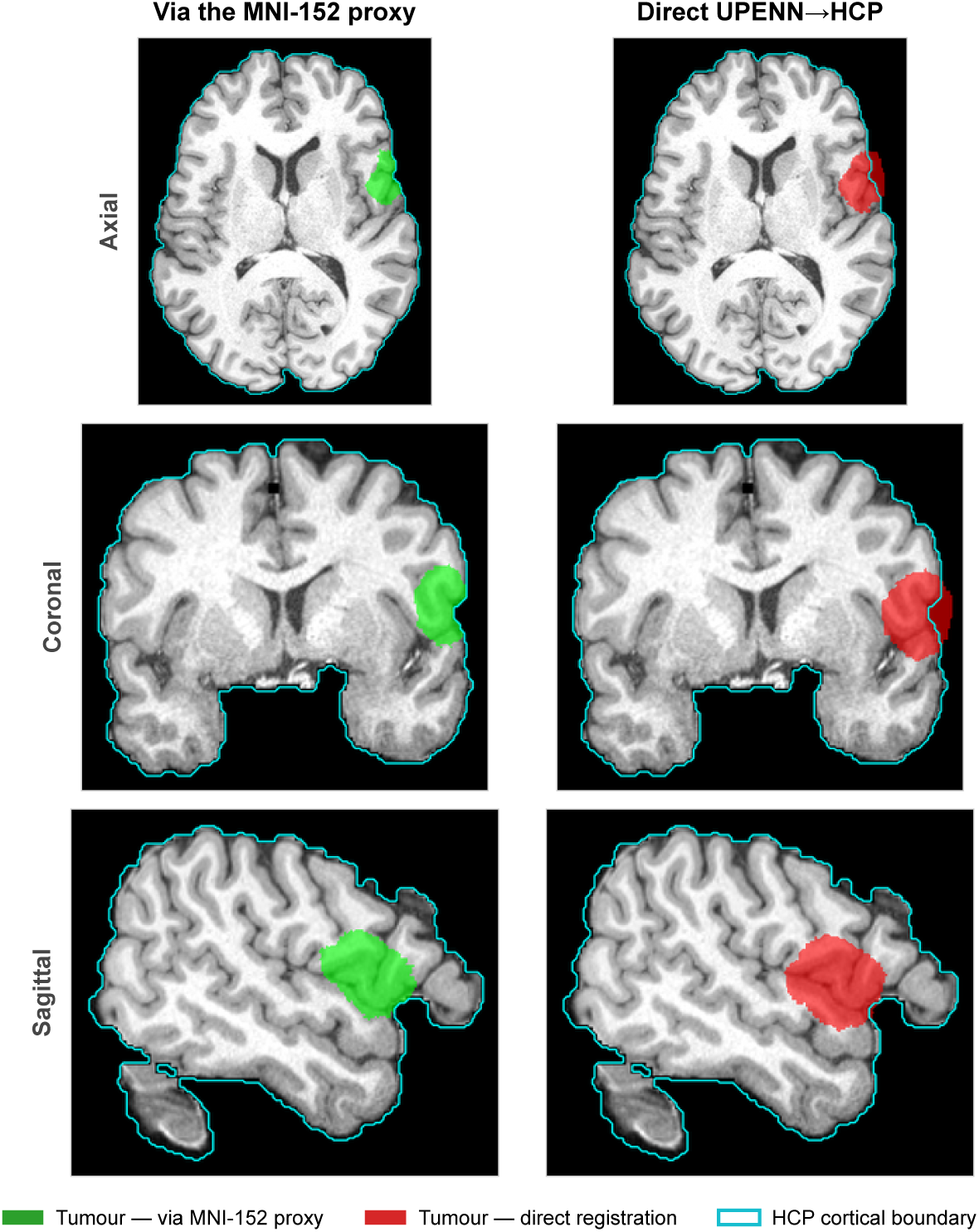
Why the tumour is transferred through the MNI152 proxy, shown for one example (UPENN-GBM tumour 20 on an exemplar HCP host) in three orthogonal views. The patient tumour mask is carried into host space either through the MNI152 template (left; the patientto-MNI and inverse host-to-MNI transforms are composed, and a tumour-exclusion mask keeps the lesion from biasing the patient-to-MNI alignment) or by direct UPENN*→*HCP registration (right). The cyan contour is the HCP cortical boundary. Direct registration pushes part of the lesion outside the cortex (16.9% of the transferred tumour in this example), whereas the MNI proxy keeps it within the brain (0.0%); nearest-neighbour interpolation maps each transferred voxel unambiguously to the original segmentation.

Registration used Advanced Normalization Tools (ANTs, version 2.5.3) (Avants et al., 2008, 2011), with all inputs reoriented to canonical RAS+ via nibabel (Brett et al., 2020). A tumour-exclusion mask (whole-brain UPENN mask minus tumour) was the moving mask for the UPENN-to-MNI registration, so tumour voxels did not contribute to the similarity metric. The UPENN T1 was registered to MNI with a composite Rigid, Affine, and nonlinear Symmetric Normalization (SyN) transform (Avants et al., 2008; Klein et al., 2009), using mutual information for the Rigid and Affine stages for robustness to inter-scanner intensity differences (Wells et al., 1996) and cross-correlation for SyN, on a hierarchical schedule of shrink factors 8 *×* 4 *×* 2 and smoothing 3 *×* 2 *×* 1 voxels. The same three-stage transform registered each HCP T1 to MNI. The tumour mask was carried into HCP space with antsApplyTransforms, composing (HCP *→* MNI)*^−^*^1^ *◦* (UPENN *→* MNI) (applied right to left, so a tumour voxel is mapped UPENN*→*MNI and then MNI*→*HCP), using nearest-neighbour interpolation throughout for binary labels. SyN was used for the nonlinear component because it is diffeomorphic, so its warps are topology-preserving and invertible by construction (Avants et al., 2008), and ranks among the most accurate nonlinear brain-registration algorithms across benchmarks (Klein et al., 2009).

### 2.3 dODF reconstruction

A dODF describes the angular distribution of water diffusion within each voxel and can represent multiple candidate directions that a single diffusion tensor cannot (Tuch, 2004; Aganj et al., 2010). Figure 2 shows examples of anisotropic and near-isotropic dODFs. dODFs were computed per HCP subject with DIPY (version 1.9) (Garyfallidis et al., 2014). The diffusion volume and gradient tables were reoriented to RAS+, the *b*-vectors rotated consistently and renormalised for non-zero *b*-values, and a diffusion-weighted brain mask restricted all voxel-wise computation. A Simple Harmonic Oscillator-based Reconstruction and Estimation (SHORE) model (Őzarslan et al., 2009; Merlet and Deriche, 2013), which gives a continuous functional basis across multi-shell data and generalises q-ball imaging to the multi-shell case, was fit voxel-wise with radial order 6 and Laplacian regularisation (*λ_N_* = *λ_L_* = 10*^−^*^8^). The dODF was sampled on the DIPY repulsion724 sphere of 724 approximately uniform directions, then converted to spherical-harmonics coefficients with the MRtrix3 command amp2sh at *l*_max_ = 12 (Tournier et al., 2019); the even-only basis at *l*_max_ = 12 gives 91 real coefficients per voxel. The spherical-harmonic coefficient volume was resampled with mrtransform -reorient fod yes, using MRtrix3’s apodised point-spread-function reorientation method, which was developed for fibre orientation distributions (Raffelt et al., 2012; Tournier et al., 2019). In the present pipeline, the same spherical-harmonic reorientation was applied to the dODF angular field. The dODF was chosen deliberately rather than a fibre orientation distribution (FOD). An FOD estimates the orientations of underlying fibre populations and is commonly used as an input to anatomical tractography (Tournier et al., 2007). Here, the aim was not to reconstruct white-matter tracts, but to use the angular distribution of water diffusion as a directional prior on simulated tumour propagation. The dODF was therefore supplied to iFOD2 as the host-specific angular field.

**Figure 2:**
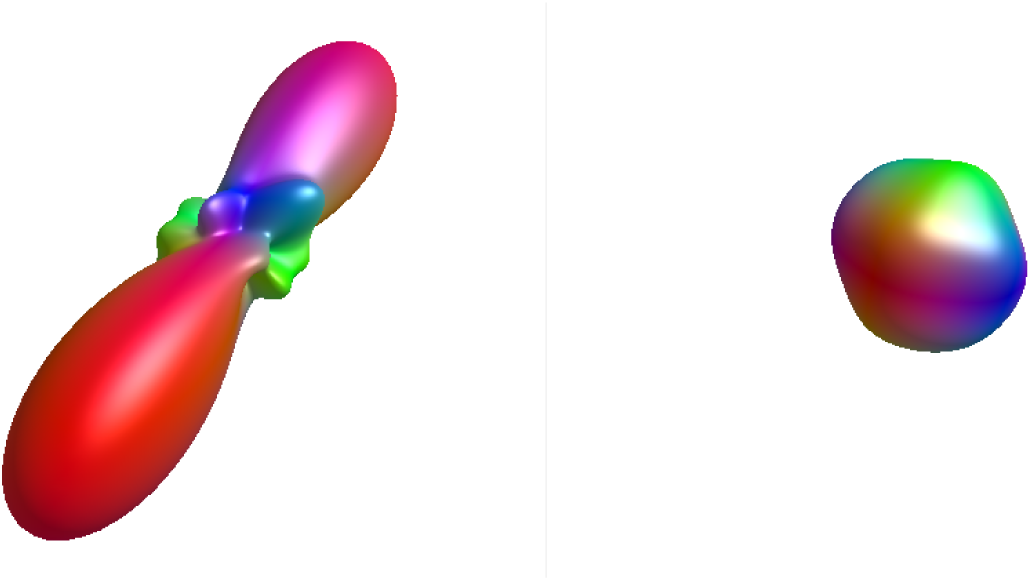
Diffusion orientation distribution functions. Left: an anisotropic voxel with a single dominant lobe marking a clear preferred direction. Right: a near-spherical isotropic voxel with no dominant direction. The simulator biases streamline propagation toward the dominant lobe or lobes of the local dODF.

### 2.4 Streamline growth simulator

The simulator is built on the MRtrix3 streamline generator tckgen (Tournier et al., 2019) using the second-order integration over fibre orientation distributions algorithm (iFOD2) (Tournier et al., 2010), which propagates a curve through a voxelised directional field by sampling candidate steps weighted by local amplitude. Although iFOD2 was developed for integration over fibre orientation distributions, the implementation accepts an antipodally symmetric spherical amplitude field represented in the MRtrix spherical-harmonic basis. In BRIAN, that field is a diffusion ODF rather than an estimate of fibre populations. The SHORE diffusion ODF is supplied in that representation, and iFOD2 is used as a second-order stochastic integrator over the diffusion-derived angular prior; the resulting streamlines are therefore interpreted as simulated propagation trajectories rather than anatomical white-matter tractography. Each streamline is one realisation of a directionally biased random walk along which tumour cells could plausibly progress given the local dODF prior, and the ensemble is aggregated into a spatial occupancy map treated as the simulated tumour. Higher-order biological processes such as proliferation, necrosis, and mass effect are not modelled.

Four parameter families are searched: a maximum turning angle angle in degrees, which limits local streamline curvature; a maximum streamline length maxlen in millimetres, a length budget for the spatial extent of invasion; a minimum-amplitude cutoff, the minimum dODF amplitude required for a step, which sharpens directional selectivity; and a three-dimensional seed coordinate (*x, y, z*), the centre of a 3 mm-radius seed sphere (the -seed sphere option), searched jointly with the others. Because these parameters are fitted to a single static mask, with no longitudinal growth or histology entering the fit, their biological labels describe how each shapes the simulation and should not be read as direct measurements of tumour biology. The integration step was left at the MRtrix3 iFOD2 default of half the voxel edge, giving 0.625 mm on the 1.25 mm dODF grid. The minimum streamline length was set to 0. A concentric exclusion sphere, one millimetre smaller in radius than the seed sphere and sharing its centre (2 mm radius against the 3 mm seed), was applied through the -exclude option, discarding streamlines that entered the seed core. Because each streamline represents a possible tumour-cell invasion path, and the seed core lies in tissue already occupied by tumour, simulated propagation does not reenter it. The excluded core is left unpopulated in the streamline-density map, and the binarised mask does not cover the tumour centre. This excluded region is small relative to the lesions, so its effect on the reported metrics is minor. Each objective evaluation inside the optimiser generated 10 000 streamlines (-select) to limit computational cost; the selected parameter vector was then re-run at 50 000, the count also used for every validation simulation, so that all reported masks derive from a denser ensemble, which resolves the normalised density field, and hence the position of the thresholded boundary, more finely.

For every configuration, tckmap voxelised the streamlines onto the HCP T1 grid with each voxel value proportional to the number of streamlines through it. The map was normalised to its maximum, giving a field in [0, 1], and a threshold of 2% of the maximum produced the binary mask. Max-normalisation removes the absolute scale of the streamline-density map, and a low threshold of 2% suppresses the diffuse, low-density stochastic tail of the streamline map; it is applied uniformly across all runs.

### 2.5 Parameter optimisation

Parameters are searched by a gradient-based procedure. Because the simulator’s output is not an analytic function of its parameters, the gradient of the objective cannot be written in closed form; it is instead estimated numerically by finite differences. Each scalar parameter is perturbed by a small amount *h* in both directions, the objective is re-evaluated at *θ* +*h* and *θ −h*, and the local slope (*f* (*θ* + *h*) *− f* (*θ − h*))*/*(2*h*) is taken as the gradient estimate that indicates which way to move that parameter. For the three-dimensional seed coordinate, simultaneous perturbation stochastic approximation (SPSA) (Spall, 1992) obtains all three components from two evaluations, drawing a random sign vector *±*1 in *x*, *y*, *z* and evaluating at *±***Δ**. The step magnitude along each estimated gradient is set by the Adam optimiser (adaptive moment estimation) (Kingma and Ba, 2015) with parameter-specific learning rates.

Per-parameter perturbation sizes, Adam learning rates, and bounds, obtained from a oneat-a-time sensitivity sweep with the other parameters at their midpoints, are listed in Table 1. Scalar parameters were seeded at the midpoint of each range (angle = 45*^◦^*, maxlen = 45 mm, cutoff = 0.075). The perturbation *h* adapts across passes, increasing when large improvements are observed and decreasing when progress stalls; after each Adam update the proposed value is projected onto the admissible bounds. For the seed coordinate, the SPSA step is followed by a short line search along the more promising direction at scales 1.5*×*, 2.0*×* and 3.0*×*. To reduce the risk of a poor local minimisation, the optimiser used a multi-seed initialisation. The binarised reference was projected onto its three principal axes, obtained by principal component analysis (PCA) of the foreground voxel coordinates, and candidates were taken at percentiles (0, 25, 50, 75, 100) along each axis. With the centre of mass, this gave 5 *×* 3 + 1 = 16 anchors, de-duplicated and reduced to the ten most widely spread by farthest-point greedy selection (Eldar et al., 1997). Each was evaluated with the scalar parameters at their midpoints, and the best-scoring candidate started the main loop. The position search was confined to a *±*5 mm window around the tumour centre of mass. The train/validation split is over hosts: for each of the 20 training hosts the optimiser runs to convergence and produces one best parameter vector, and the per-tumour component-wise median across these 20 vectors is applied once to each of the 65 validation hosts.

**Table 1:** Per-parameter bounds, initial perturbations, and Adam learning rates used by the iterative optimiser. Bounds for the three coordinate components are defined relative to each subject’s tumour centre of mass (COM) and are given in millimetres around that reference point

| Parameter | Units | Bounds | Perturbation | Adam learning rate |
| --- | --- | --- | --- | --- |
| **angle** | ° | [20, 90] | 2.0 | 10.0 |
| **maxlen** | mm | [20, 90] | 2.0 | 10.0 |
| **cutoff** | dimensionless | [0.04, 0.10] | 0.005 | 0.01 |
| *x* (relative to COM) | mm | ±5 | 1.0 | 1.0 |
| *y* (relative to COM) | mm | ±5 | 1.0 | 1.0 |
| *z* (relative to COM) | mm | ±5 | 1.0 | 1.0 |

### 2.6 Evaluation metrics and constrained objective

During per-host optimisation and validation, the reference tumour mask was modified by excluding a small central neighbourhood around the seed position, approximately corresponding to the exclusion region used during streamline generation. By contrast, all aggregate-mask analyses in MNI space were evaluated against the complete MNI-warped tumour-core mask.

Performance was evaluated using complementary overlap, boundary, surface, volume, and projection metrics, since no single measure captures all aspects of spatial agreement (Maier-Hein et al., 2024; Taha and Hanbury, 2015). Overlap was quantified by the Dice coefficient (DSC),

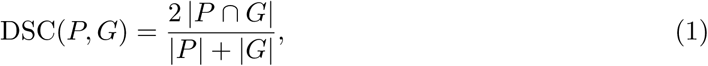

expressed as a loss dice loss = 1 *−* DSC(*P, G*). Boundary error was measured using a 95thpercentile Hausdorff distance (Taha and Hanbury, 2015). In accordance with the implementation, directed nearest-surface distances were calculated in both directions, pooled, and reduced to their 95th percentile:

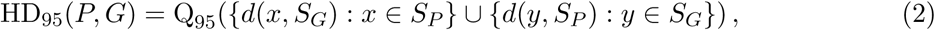

where *S_P_* and *S_G_* are the surface voxels of the simulated and reference masks, respectively, *d* is the nearest-surface distance, and Q_95_ denotes the 95th percentile. Distances were calculated in millimetres using the voxel spacing recorded in the image header. For use in the optimisation objective, HD_95_ was converted to a bounded loss term, hd norm = min(HD_95_*/*HD DMAX MM, 1.0), with HD DMAX MM = 20 mm. The 20-mm cap was a fixed parameter of the present implementation. Surface Dice within a tolerance *τ* (Nikolov et al., 2021),

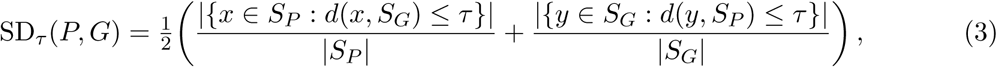

uses *τ* = SURF TOL MM = 2 mm, with sd loss = 1 *−* SD*_τ_* . A relative volume-error term,

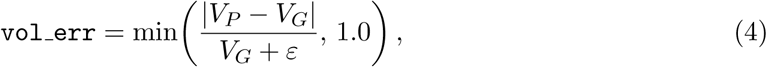

with *ε* = 10*^−^*^10^, makes size mismatch explicit. Finally, three silhouette projections by logical OR along each orthogonal axis give projection-Dice losses

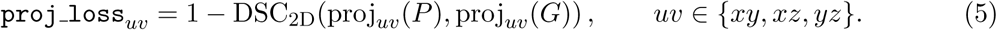

These are aggregated as the worst case

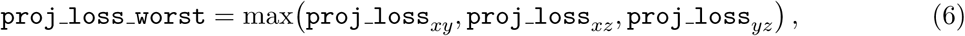

so two correct projections cannot mask a severe mismatch in the third.

The base objective is a weighted linear form,

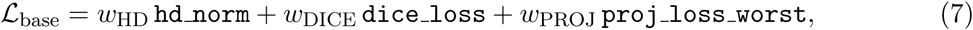

with stage-dependent weights summing to one. Surface-Dice and volume error enter as feasibility constraints: a mask is feasible when dice loss *≤ ε*_dice_ *∧*sd loss *≤ ε*_sd_ *∧*vol err *≤ ε*_vol_. Outside the feasible region, constraint violations are quantified as

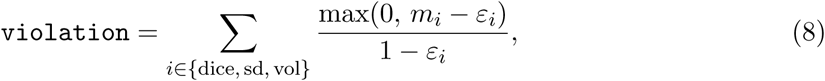

The constraints were incorporated using a penalty formulation (Nocedal and Wright, 2006). The final objective augments the base objective with the resulting penalty,

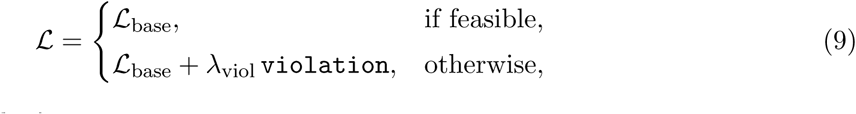

where the penalty weight *λ*_viol_ was set to 2.0.

The objective is tightened through a three-stage curriculum. Stage A gives the boundary term the largest weight, (*w*_HD_*, w*_DICE_*, w*_PROJ_) = (0.55, 0.45, 0.00) with (*ε*_dice_*, ε*_sd_*, ε*_vol_) = (0.45, 0.55, 0.15), placing the surface roughly where the reference surface lies. Each promotion multiplies *ε*_sd_ and *ε*_vol_ by 0.9 while *ε*_dice_ is held fixed, giving stage B weights (0.35, 0.55, 0.10) with (0.45, 0.495, 0.135) and stage C weights (0.20, 0.60, 0.20) with (0.45, 0.446, 0.122). A streak rule promotes after *k* consecutive feasible evaluations (*k* = 5 for A*→*B, *k* = 8 for B*→*C); a patience rule promotes when the stage’s best objective fails to improve by more than improve tol = 0.005 for patience passes = 10 consecutive passes. The optimiser terminates when stage C exhausts its patience budget or the running best objective falls below target obj = 0.1.

### 2.7 Aggregation and the directional-prior comparisons

Aggregation was performed in MNI space. As described above, each per-host streamline-density map was normalised to its maximum and binarised at a threshold of 0.02. The resulting binary masks were warped into MNI space and averaged voxel-wise across hosts, producing a host-frequency map for each tumour. Separate training and validation host-frequency maps were generated from 20 and 65 hosts, respectively, and each was thresholded at a host-frequency value of 0.02 to produce an aggregate binary mask. The reference mask was also warped into MNI space, and the full metric suite was recomputed on the aggregate masks. Validation masks were scored on the fixed stage-C scale, the strictest, so masks from different curriculum stages remained comparable.

The directional prior was probed on a purposive subset of 10 of the 30 tumours (UPENNGBM-00001, 00019, 00020, 00023, 00024, 00025, 00031, 00032, 00034, and 00042) chosen to span the cohort’s performance and size range, including high-, mid-, and low-accuracy cases. The subset’s per-tumour median Dice (0.757) was close to that of the full cohort (0.748), while its wider spread reflected the deliberate inclusion of both lower- and higher-performing cases. The parameter bounds (Table 1) were derived from pilot tuning on a single tumour, so the largest and smallest lesions are expected to fit less well, but a fixed bound range is necessary to keep the search tractable and to stop it degenerating toward a grid search, and the subset deliberately includes such cases.

Three tracking conditions were run on this subset, all on the identical 20 training and 65 validation hosts and the full three-stage curriculum, and all operating on an orientation distribution function, so that any difference between conditions is attributable to the diffusion model or the tracker alone. The dODF condition is the method described above, a SHORE dODF reconstructed from all shells and propagated with iFOD2. The isotropic null replaces only the tracker, running the MRtrix3 tckgen NullDist2 algorithm, the null-distribution counterpart of iFOD2 that samples propagation directions at random subject to the same curvature and length constraints (Morris et al., 2008; Tournier et al., 2019), on the identical SHORE dODF file, so it is the direction-blind floor of the same pipeline. The tensor-dODF condition replaces only the diffusion model, fitting a single diffusion tensor from the *b ≤* 1000 s/mm^2^ shell with DIPY (Basser et al., 1994; Garyfallidis et al., 2014) and sampling its diffusion ODF on the same repulsion724 sphere, converted with the same amp2sh order and reoriented onto the same grid as the SHORE dODF; it is therefore a drop-in prior tracked with identical iFOD2 settings, carrying one ellipsoidal lobe per voxel where the SHORE dODF can carry several (Figure 3). The SHORE dODF was compared with the isotropic null to assess the effect of directional sampling, and with the tensor dODF to assess whether resolving multiple intravoxel orientations improved performance.

**Figure 3:**
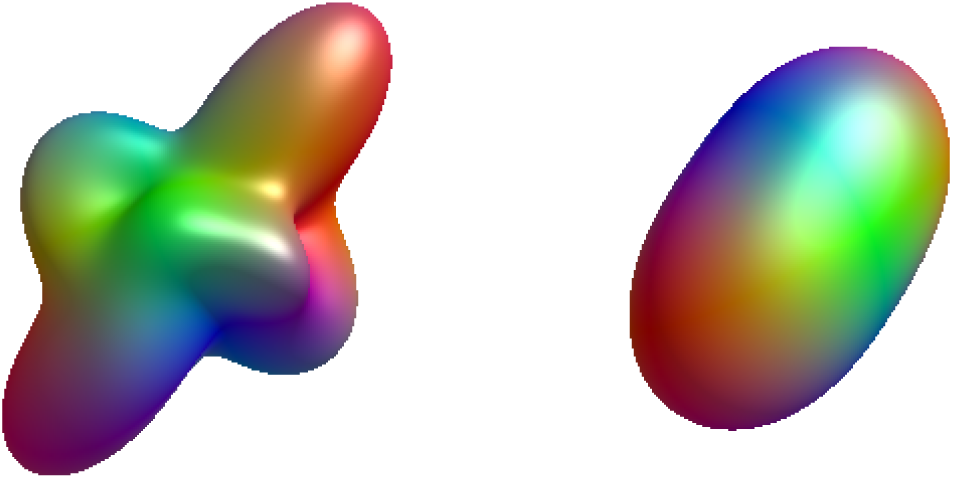
Resolving crossing fibres. Left: the SHORE dODF resolves the multiple fibre populations at a crossing as distinct lobes. Right: a single diffusion tensor from the same voxel, recast as a dODF, collapses the crossing into one ellipsoidal lobe with a single dominant orientation. The tensor-dODF ablation condition tests whether resolving these crossings improves the directional prior over a single-fibre model.

**Figure 4:**
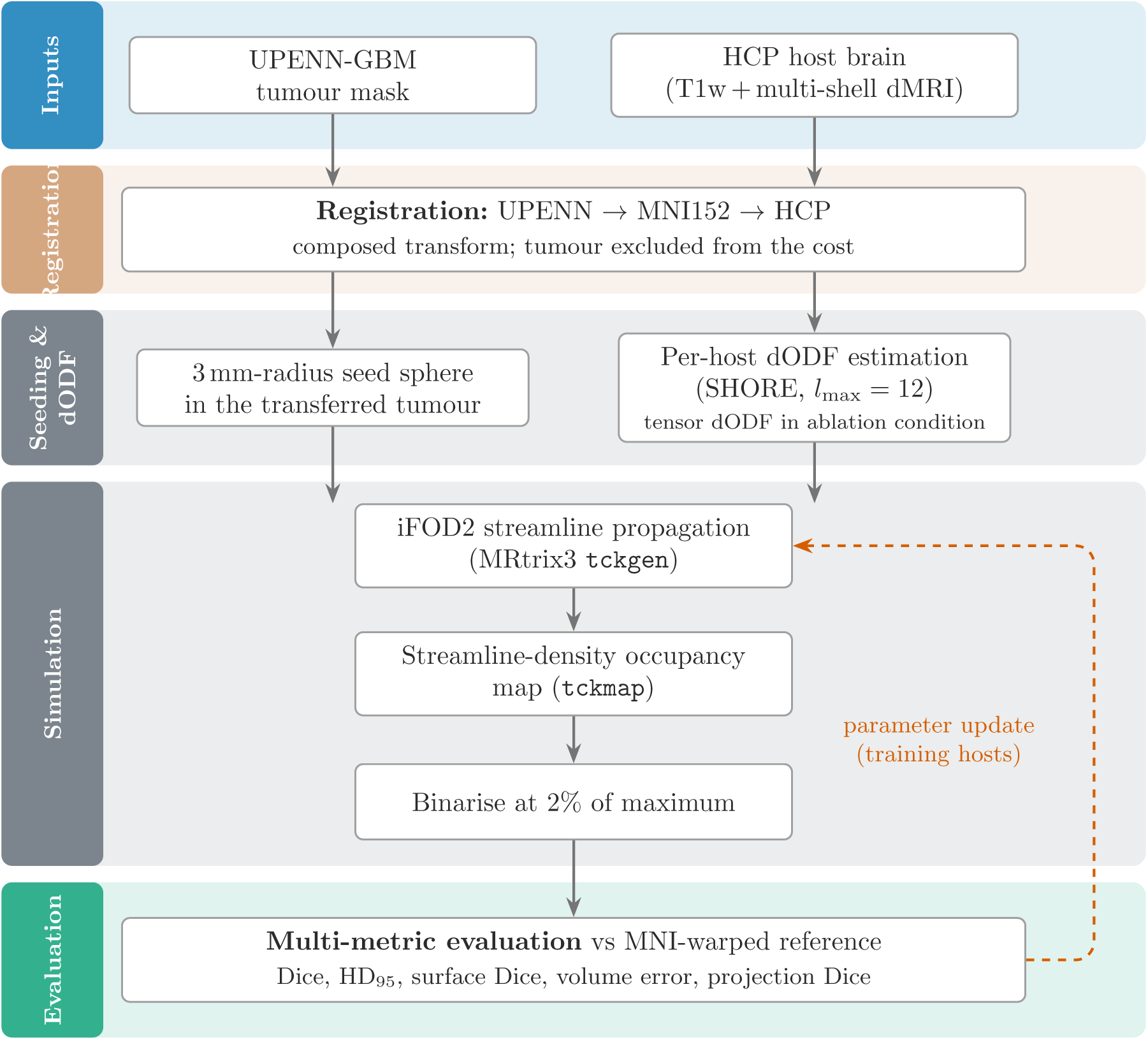
Overview of the BRIAN pipeline. A patient tumour mask (UPENN-GBM) and a healthy host brain (HCP) are each registered to the MNI152 template; composing the two transforms, one inverted, carries the tumour mask into host space without forcing the host brain to match the tumour region. A dODF is reconstructed on the host, stochastic iFOD2 streamlines are seeded within the transferred mask and propagated under the dODF prior, and the streamline-density map is thresholded into a binary simulated mask that is compared against the patient reference. The optimiser updates the propagation parameters on the 20 training hosts; the 65 validation hosts are simulated once with the median parameters. In the directional-prior ablation (Section 3.5) the SHORE dODF of the seeding-and-dODF step is replaced by a single-tensor dODF or by a direction-blind null, with all other stages unchanged.

For each condition, the 65 per-host binary masks were averaged voxel-wise to produce a host-frequency map for each tumour. This map was thresholded at a host-frequency value of 0.02 and scored once against the reference, giving one value per tumour per condition on every metric. Inference was therefore conducted at the tumour level with no nested pairs: for each contrast, the 10 per-tumour differences were tested against zero using a one-sample *t*-test (*n* = 10), Holm-corrected across the five metrics, with a Wilcoxon signed-rank test on the same 10 values as a non-parametric check. Results are reported using the mean difference, its 95% confidence interval, and Cohen’s paired effect size *d_z_*. Scoring one aggregate mask per tumour avoids treating individual host runs as independent tumour-level observations.

### 2.8 Implementation

The pipeline was implemented under Linux with Python 3.12, numpy 1.26+ (Harris et al., 2020), scipy 1.15 (Virtanen et al., 2020), scikit-learn 1.4 (Pedregosa et al., 2011), pandas 2.2 (McKinney, 2010), and nibabel 5.3 (Brett et al., 2020). Neuroimaging operations are delegated to MRtrix3 3.0.4 (Tournier et al., 2019) for diffusion reconstruction, streamline generation, and volumetric conversion, to ANTs 2.5.3 (Avants et al., 2008, 2011) for registration, and to DIPY 1.9 (Garyfallidis et al., 2014) for SHORE fitting, sphere sampling, and spherical-harmonics conversion. The full source code is available in the public repository accompanying this article.

## 3 Results

The evaluation comprised 30 UPENN-GBM patient tumour masks, each propagated onto 65 HCP validation hosts, for 1 950 tumour-host runs. Five metrics are reported per run, all oriented so that higher values indicate closer agreement with the patient reference (Maier-Hein et al., 2024): the Dice coefficient, a bounded Hausdorff agreement 1*−*min(HD_95_*/*20 mm, 1), the surface Dice at a 2 mm tolerance, a bounded volume match 1 *−* min(*|*Δ*V |/V*_GT_, 1), and the worst-case projection Dice over the three orthogonal projections.

### 3.1 Cohort-level performance

Pooled across all 1 950 pairs, the simulator produced a median Dice of 0.745 (IQR [0.688, 0.782]); this is the pooled per-run estimator. The median Hausdorff agreement was 0.776 (median HD_95_ approximately 4.48 mm in MNI space), the median surface Dice at 2 mm 0.662, the median volume match 0.798, and the median worst-case projection Dice 0.824. Figure 5 shows the cohort-level distribution of each metric. Dice, surface Dice, and worst-projection Dice had similar medians. Worst-projection Dice showed the narrowest distribution, whereas volume match showed the widest IQR.

**Figure 5:**
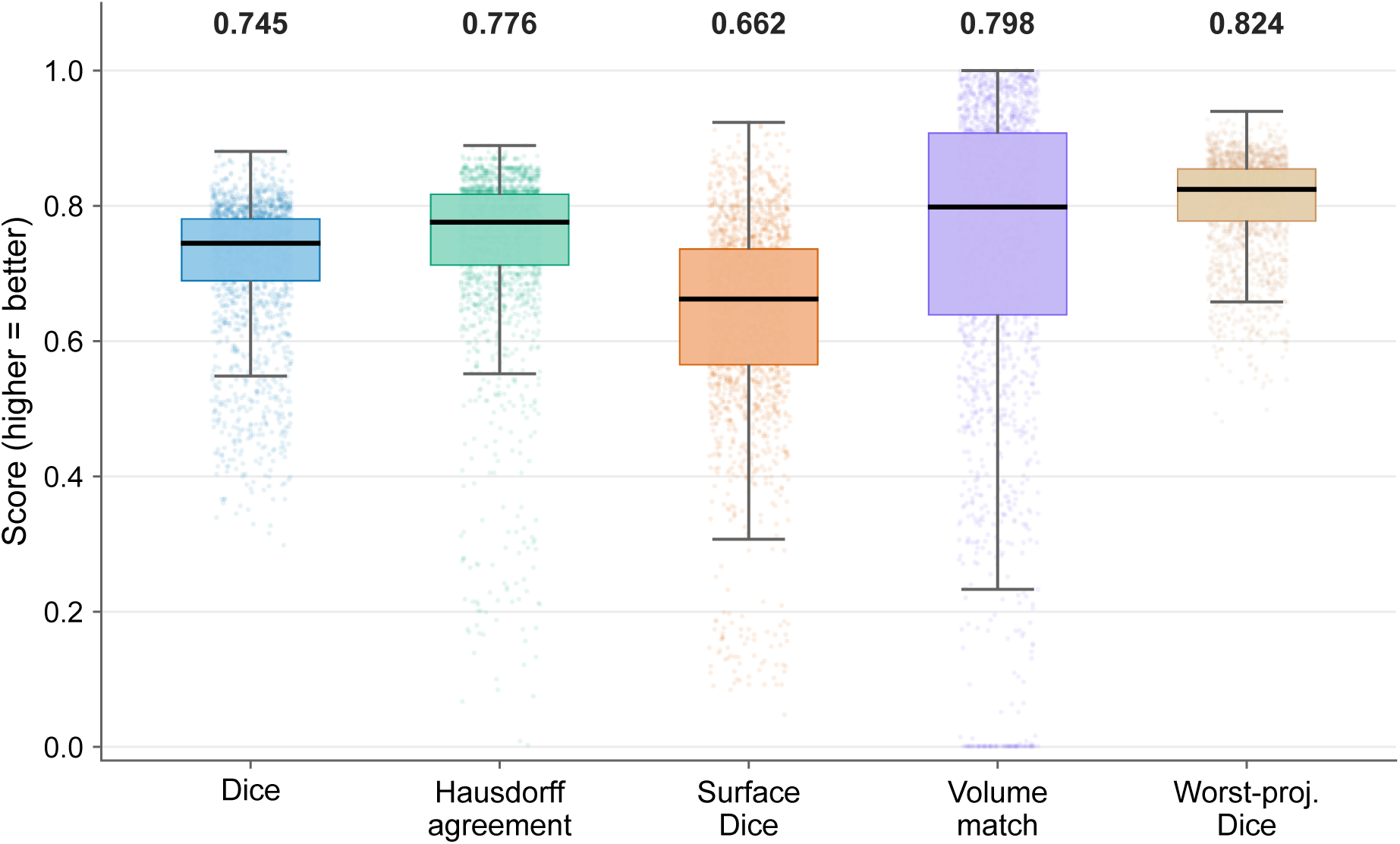
Cohort-level distribution of the five evaluation metrics across the 1 950 validation tumour-host pairings (30 tumours *×* 65 hosts). Boxes mark the inter-quartile range, whiskers the 1.5*×*IQR envelope, and points the outliers; the cohort median is printed above each box. All metrics are oriented so that higher values are better; Hausdorff agreement is the bounded form 1 *−* min(HD_95_*/*20 mm, 1) and volume match 1 *−* min(*|*Δ*V |/V*_GT_, 1).

### 3.2 Per-tumour profile and host reproducibility

The median of the 30 per-tumour median Dice values was 0.748 (IQR [0.704, 0.777]), close to the pooled per-run median of 0.745. 23 of the 30 tumours produced a per-tumour median Dice above 0.70, and 13 exceeded 0.75. The best tumour, UPENN-GBM-00035, reached a per-tumour median Dice of 0.844, with a median Hausdorff agreement of 0.844, surface Dice of 0.840, volume match of 0.910, and worst-projection Dice of 0.895. The lowest-tail tumours were UPENN-GBM00001 (0.419), UPENN-GBM-00031 (0.479), UPENN-GBM-00028 (0.509), UPENN-GBM-00024 (0.604), and UPENN-GBM-00005 (0.615), with UPENN-GBM-00018 (0.643) close behind. Visual inspection suggested that lower-performing cases were more common at the smallest and largest tumour volumes, whereas higher Dice values were more frequent among mid-sized lesions. However, there was no clear monotonic association between reference tumour volume and median Dice (Spearman *ρ* = 0.07; Figure 6). Across tumours, Dice IQRs were generally only a few hundredths and did not exceed about

**Figure 6:**
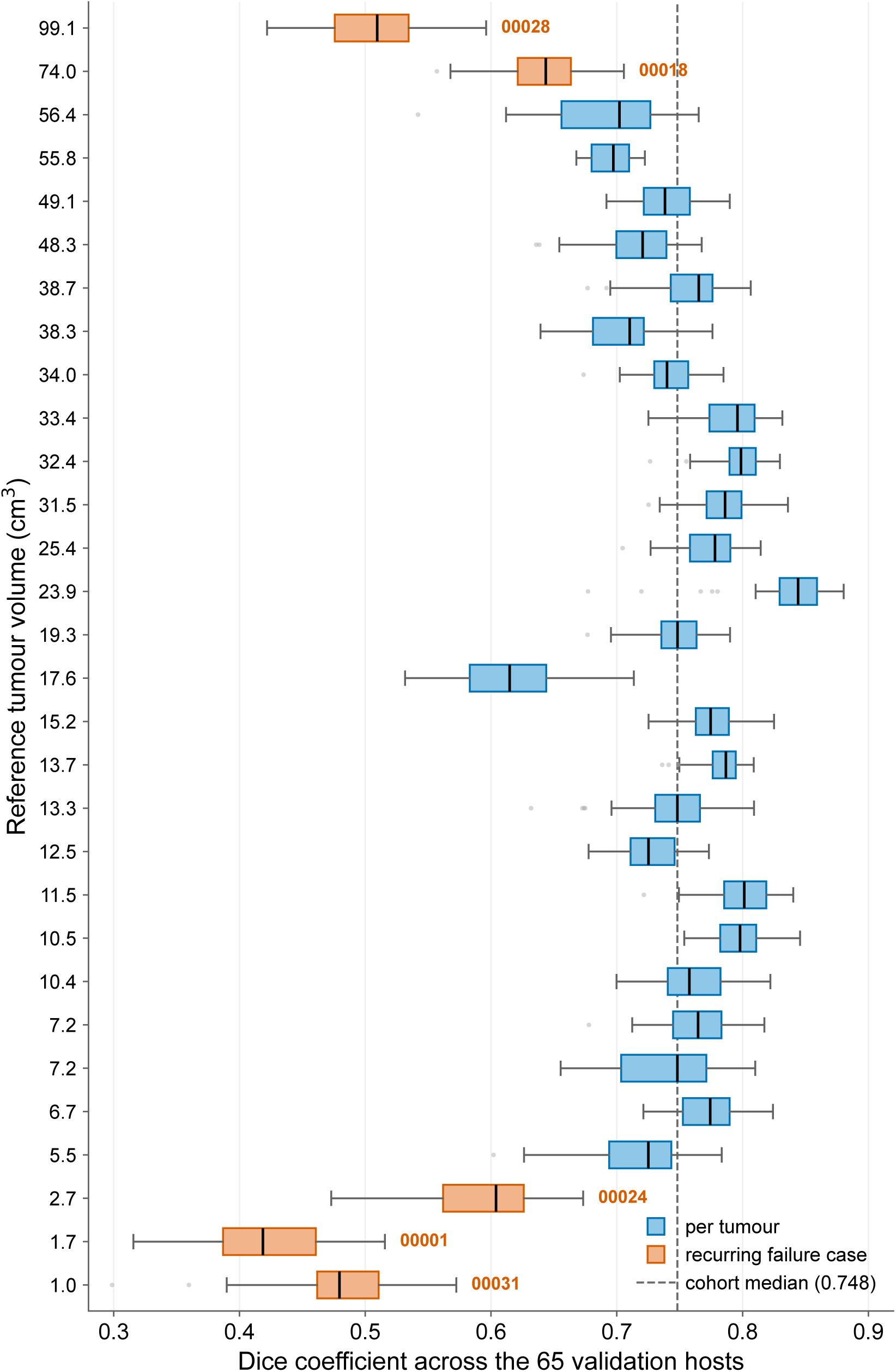
Per-tumour distribution of the Dice coefficient across the 65 validation hosts, one box per tumour, ordered by reference tumour volume (smallest at the bottom). The dashed line marks the cohort median. Boxes mark the inter-quartile range and whiskers the 1.5*×*IQR envelope; the recurring low-accuracy cases of Section 3.4 sit at the volume extremes. 0.08. The other placement and shape metrics showed similarly limited variation across hosts, although volume match was less stable in a small number of runs. Figure 7 shows the validation host-frequency contours for six tumours spanning the observed performance range.

### 3.3 Averaged masks in MNI space

For each tumour, the binary per-host masks were averaged voxel-wise separately across the 20 training hosts and the 65 validation hosts. Each resulting host-frequency map was thresholded at 0.02 and scored against the MNI-warped patient reference. The median Dice of the validation aggregate masks was 0.707, compared with 0.639 for the training aggregate masks. Hausdorff agreement was also higher for the validation aggregates than for the training aggregates (0.646 vs. 0.523), as was volume match (0.267 vs. 0.000). The validation-average exceeded the training-average on Dice in 28 of 30 tumours and on Hausdorff agreement in 27 of 30; the two exceptions on Dice were UPENN-GBM-00021 (negligible) and UPENN-GBM-00028 (training-average higher by 0.107).

These aggregate-mask values were computed using a host-frequency threshold of 0.02 and are not comparable to the per-host medians above or to the 0.5 display threshold used in Figure 7. Because the training and validation aggregates contain different numbers of hosts, comparisons between them are descriptive.

**Figure 7:**
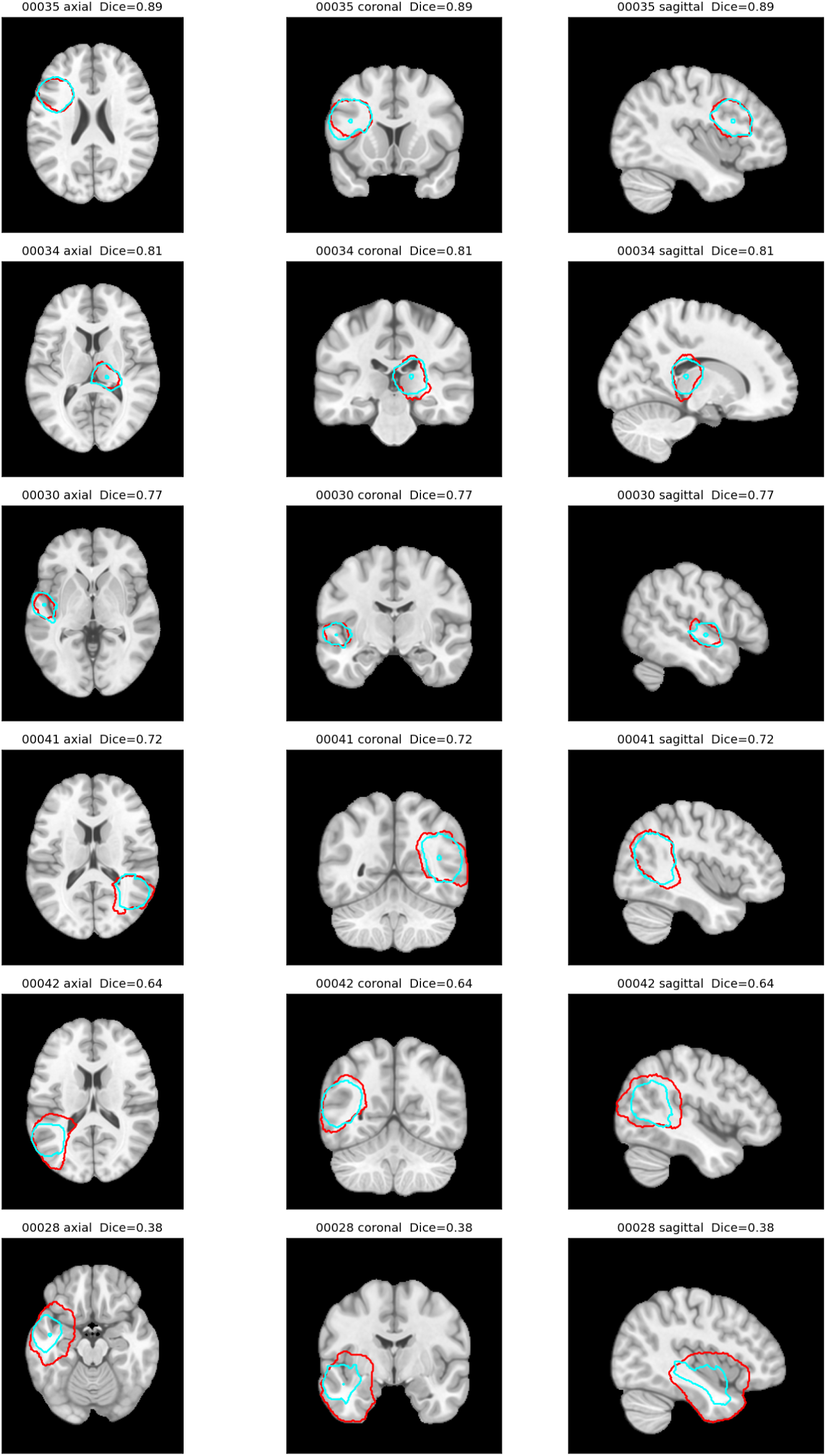
Orthogonal (axial, coronal, sagittal) montage in MNI space for six representative tumours ordered top to bottom from best to worst by validation-average Dice. The red contour is the patient-specific ground truth; the cyan contour is the validation host-frequency map thresholded at 0.5, so a voxel is displayed when it is present in at least half of the 65 validation hosts.

### 3.4 Recurring error patterns

Two recurring error patterns accounted for most of the lower tail. The first is a small or peripheral lesion with a depressed or floored volume match. UPENN-GBM-00001, a small right cerebellar lesion, had a median Dice of 0.419 and a median volume match of exactly zero (the volume-error term saturating across all 65 hosts), while its Hausdorff agreement stayed comparatively high at 0.736, surface Dice 0.561, and worst-projection Dice 0.610: a mask correctly placed but systematically larger than the patient lesion. The tendency recurred in milder form on UPENN-GBM-00031 (median Dice 0.479, Hausdorff agreement 0.774, surface Dice 0.640, volume match 0.233) and UPENN-GBM-00024 (median Dice 0.604, Hausdorff agreement 0.818, surface Dice 0.701, volume match 0.614); only UPENN-GBM-00001 reached the volume-match floor across all 65 hosts. The placement-sensitive metrics stayed interpretable, consistent with reporting boundary- and volume-sensitive metrics alongside overlap (Maier-Hein et al., 2024).

The second pattern is a large lesion with systematic under-extension. UPENN-GBM-00018, an extensive right-temporal lesion, had a median Dice of 0.643, Hausdorff agreement of 0.232 (median HD_95_ approximately 15.4 mm), surface Dice of 0.456, volume match of 0.511, and worst-projection Dice of 0.718, with the simulated contour concentric but uniformly smaller in all three projections. This was more severe on UPENN-GBM-00028: median Dice 0.509, Hausdorff agreement 0.510 (median HD_95_ approximately 9.8 mm), surface Dice 0.146, volume match 0.344, and worst-projection Dice 0.685. The first pattern occurs when the fixed global threshold is applied to a smaller-than-typical lesion, and the second when the fixed streamline budget is applied to a larger-than-typical one.

### 3.5 Isolating the directional prior: null, SHORE-dODF, and tensor-dODF conditions

Three conditions were compared on the purposive ten-tumour subset, each aggregated to a single validation aggregate mask per tumour so that inference is at the tumour level (*n* = 10, Section 2.7): the SHORE dODF condition, an isotropic null that shares the dODF file but tracks direction-blind, and a tensor-dODF condition that swaps the SHORE model for a single tensor under the same tracker. Table 2 reports the condition means and the dODF-versus-null contrasts, and Figure 8 shows the per-tumour Dice.

**Figure 8:**
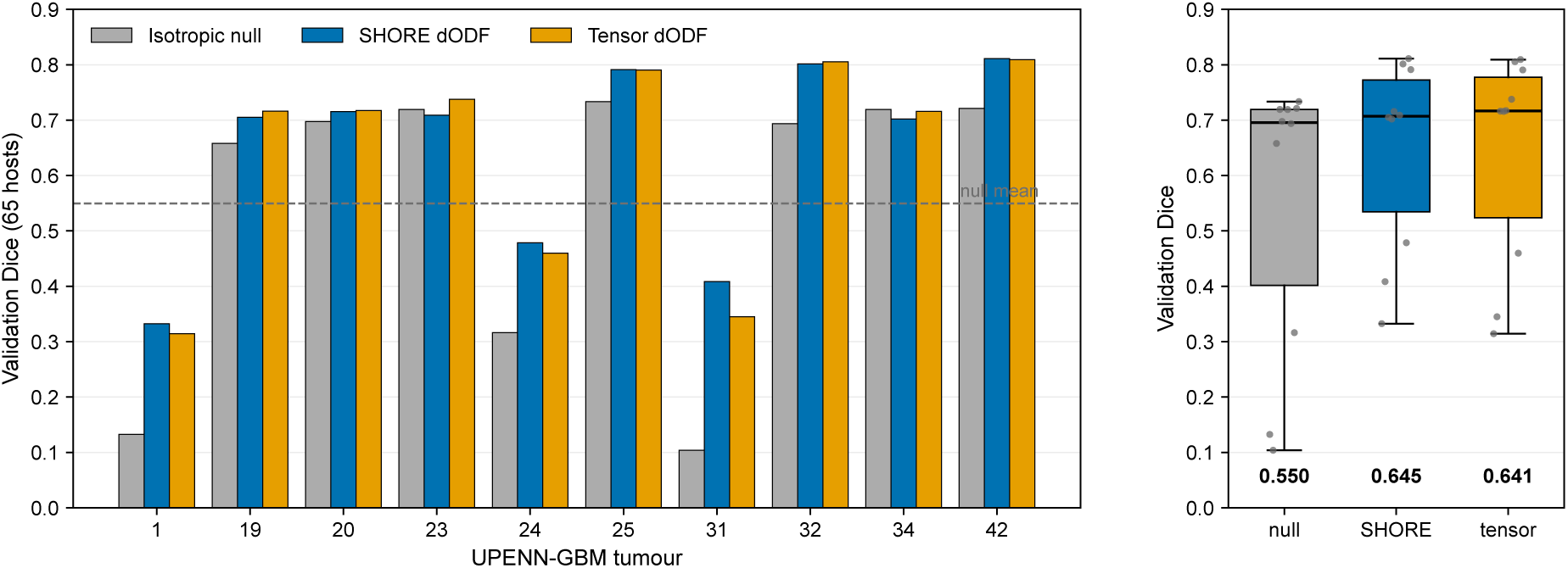
Validation Dice for the three tracking conditions on the 10-tumour subset (aggregate mask over 65 held-out hosts per tumour). Left: per-tumour Dice for the isotropic null (NullDist2), the SHORE dODF, and the tensor dODF (both iFOD2); the dashed line marks the null mean. Right: the distribution across the ten tumours, with the condition mean printed below each box. Both dODF conditions outperformed the null on most tumours, and no statistical difference was observed between the two conditions; see Table 2.

**Table 2:** Directional-prior ablation on the 10-tumour subset (validation aggregate mask, one value per tumour, *n* = 10). **Top:** mean of each metric for the three conditions, all sharing the seeding, curvature, length, and streamline budget; the isotropic null tracks the SHORE dODF direction-blind (NullDist2), and the two dODF conditions track with iFOD2 and differ only in the diffusion model. Higher is better on every column (Hausdorff agreement is the bounded boundary score). **Bottom:** the SHORE dODF versus isotropic null contrast at the tumour level, with the mean per-tumour difference (dODF *−* null), its 95% confidence interval, Cohen’s paired *d_z_*, the Holm-corrected one-sample *t*-test across the five metrics, and the Wilcoxon signed-rank *p*. The tensor dODF gave a comparable separation from the null, and no statistically detectable difference from the SHORE dODF was observed on any metric (Section 3.5).

| Condition (tracker) | Dice | Hausdorff agr. | Surface Dice | Volume match | Worst-proj. Dice |
| --- | --- | --- | --- | --- | --- |
| Isotropic null (NullDist2) | 0.550 | 0.558 | 0.250 | 0.228 | 0.635 |
| SHORE dODF (iFOD2) | 0.645 | 0.673 | 0.401 | 0.311 | 0.725 |
| Tensor dODF (iFOD2) | 0.641 | 0.666 | 0.405 | 0.339 | 0.714 |

Table 2: Directional-prior ablation on the 10-tumour subset (validation aggregate mask, one value per tumour, $n = 10$ ).
| SHORE dODF – null | Mean $\Delta$ [95% CI] | $d_z$ | $p$ Holm; Wilcoxon |
| --- | --- | --- | --- |
| Dice | +0.096 [+0.023, +0.169] | 0.94 | 0.061; 0.010 |
| Hausdorff agreement | +0.115 [+0.009, +0.221] | 0.78 | 0.073; 0.027 |
| Surface Dice | +0.150 [+0.059, +0.241] | 1.18 | 0.023; 0.004 |
| Volume match | +0.083 [−0.053, +0.219] | 0.44 | 0.200; 0.297 |
| Worst-proj. Dice | +0.090 [+0.015, +0.165] | 0.86 | 0.071; 0.020 |

Compared with the isotropic null, the SHORE dODF increased Dice by 0.096 (*d_z_* = 0.94), surface Dice by 0.150 (*d_z_* = 1.18), worst-projection Dice by 0.090 (*d_z_* = 0.86), and Hausdorff agreement by 0.115 (*d_z_* = 0.78), each in eight to nine of ten tumours. Volume match increased by 0.083 (*d_z_*= 0.44) in four of ten tumours. All five mean differences were positive, but after Holm correction across the five metrics only the surface-Dice contrast remained significant (Table 2). The tensor-dODF condition showed comparable differences from the null (*d_z_* up to 1.58). Directional sampling therefore improved agreement with the reference masks on every metric in this subset, though only one contrast survived correction for multiple comparisons at *n* = 10.

The SHORE and tensor dODF conditions produced similar results. Mean differences between the two conditions were below 0.03 for every metric, with all *|d_z_| ≤* 0.51 and Holm-corrected *p ≥* 0.70. This subset therefore provided no evidence that resolving multiple intravoxel orientations improved agreement with the visible tumour envelope. These values were calculated from one aggregate validation mask per tumour and should not be compared directly with the full-cohort per-host medians in Section 3.1.

## 4 Discussion

BRIAN was calibrated and evaluated against the MRI-visible tumour core, comprising enhancing tumour and necrotic core with oedema excluded. The study therefore assesses reproduction of the visible lesion rather than occult invasion beyond its margin.

The directional prior was derived from healthy HCP white matter and transferred to the patient lesion through patient-to-MNI-to-host registration. Glioblastoma can distort or destroy local tracts (Price et al., 2003; Claes et al., 2007), so the host diffusion field cannot reproduce the patient’s pre-diagnostic anatomy. It was used because pre-diagnostic diffusion MRI was not available in the source cohort.

### 4.1 Cohort performance and directional-prior comparisons

Across the 30 tumours, the per-tumour median Dice was 0.748 (IQR [0.704, 0.777]), with a pooled median of 0.745 across the 1 950 tumour–host pairings. Median Hausdorff agreement was 0.776. Parameters were not re-optimised for individual validation hosts. Since the train/validation split was defined over hosts rather than tumours, these results assess reproduction of known lesions across host anatomies rather than prediction of unseen tumours.

The isotropic NullDist2 condition sampled directions randomly while retaining the same curvature, length, and streamline-budget settings (Morris et al., 2008; Tournier et al., 2019). On the ten-tumour subset, the SHORE dODF condition improved every overlap and boundary metric relative to this null condition, in eight to nine of the ten tumours. Volume match also improved, in four of ten tumours. These results indicate that directional sampling contributed to agreement with the reference masks, although only the surface-Dice contrast remained significant after correction for multiple comparisons.

The SHORE and tensor dODF conditions produced similar metric values, with no statistically detectable differences on this subset. The present data therefore provide no evidence that resolving multiple intravoxel orientations improved agreement with the MRI-visible tumour core.

### 4.2 Relation to previous work

Previous mechanistic glioma models have used diffusion MRI to represent anisotropic invasion. Jbabdi et al. (2005) used DTI to simulate anisotropic low-grade glioma growth, while Clatz et al. (2005) incorporated white-matter diffusion information into a three-dimensional growth model with biomechanical deformation. Painter and Hillen (2013) derived a macroscopic invasion model from cell migration along DTI-defined fibre orientations, and Engwer et al. (2015) developed a related multiscale DTI-based model. Other approaches used diffusion-informed anisotropic eikonal or Riemannian formulations to model tumour propagation and invasion margins (Konukoglu et al., 2010; Mosayebi et al., 2012). Alfonso et al. (2017) reviews this broader class of models.

Patient-specific proliferation–invasion models have also been used to predict radiotherapy response (Rockne et al., 2010) and to optimise radiotherapy plans (Corwin et al., 2013); Baldock et al. (2013) reviews their potential role in precision neuro-oncology. In the present study, the tensor dODF performed similarly to the SHORE dODF. Streamline-based reconstructions are susceptible to false-positive pathways and require cautious anatomical interpretation (Jones et al., 2013; O’Donnell and Pasternak, 2015; Maier-Hein et al., 2017). They are therefore treated here as directional-plausibility priors rather than direct anatomical invasion routes.

### 4.3 Reproducibility across hosts

Variation across HCP hosts was small relative to the differences between tumours. Training and validation aggregate masks gave broadly similar results, although the comparison is descriptive because they were derived from 20 and 65 hosts, respectively.

### 4.4 Limitations

The study included 30 unifocal tumours from a single imaging cohort and was not designed to support patient-level clinical claims. Multifocal tumours were excluded by design, because the optimiser is built around a single seeded propagation, introducing a selection bias toward single-focus lesions. The IDH labels were taken from the per-subject designations in Bakas et al. (2022) without independent re-screening against the 2021 WHO criteria (Louis et al., 2021); 28 of the 30 tumours are recorded as IDH-wildtype and two as IDH-NOS/NEC (status not determined), so any upstream misclassification propagates here. The streamline-density threshold and the maximum streamline count are fixed global constants across all runs, which keeps configurations comparable but makes volume match sensitive to over-extension on small peripheral lesions and to under-extension on the most extensive ones. Validation applies a single median parameter vector per tumour to each host without per-subject re-optimisation, so the validation metrics reflect both the quality of that median vector and the homogeneity of the training distribution. Finally, the worst-projection Dice is computed on three orthogonal projections rather than the full 3-D reference, so a small systematic mis-orientation off the MNI cardinal axes goes unpenalised. The present study should therefore be interpreted as a methodological proof of concept, not as a clinically validated model of occult tumour spread.

## Data and Code Availability

The BRIAN simulator and all analysis code are openly available at https://github.com/ Aitor-Alberdi/project-brian-pipeline.

No new imaging data were generated for this study. Healthy host data are from the Human Connectome Project 1200 Subjects release, available under open-access terms via ConnectomeDB (https://db.humanconnectome.org). Glioblastoma imaging and segmentations were accessed through The Cancer Imaging Archive (TCIA) from the UPENN-GBM collection (Bakas et al., 2021; Clark et al., 2013), used under the TCIA Data Usage Policy. The HCP-to-MNI registration cache generated here can be reused by downstream studies.

## Ethics

This study used only de-identified, publicly released neuroimaging data and collected no primary data. Ethics approval was granted by the Faculty of Medicine and Surgery Research Ethics Committee of the University of Malta under Research Ethics and Data Protection (REDP) application MED-2025-00309, linked to parent project MED-2024-00356 (approved by the Faculty Research Ethics Committee and the University Research Ethics Committee Data Protection panel).

## Author Contributions

**Aitor Alberdi Escudero:** Conceptualization, Methodology, Software, Formal analysis, Investigation, Data curation, Visualization, Writing – original draft. **Kenneth Scerri:** Methodology, Supervision, Writing – review & editing. **Andrew Sammut:** Supervision, Writing – review & editing. **Claude J. Bajada:** Conceptualization, Methodology, Supervision, Writing – review & editing.

## Acknowledgements

This work is dedicated to the memory of Brian Bajada, father of Claude J. Bajada, after whom this project was named. The authors gratefully acknowledge the ALIVE Charity Foundation, a cyclist-led charity whose members raise funds through endurance cycling challenges, for their effort and commitment to supporting biomedical research. Data were provided in part by the Human Connectome Project, WU-Minn Consortium (Principal Investigators: David Van Essen and Kamil Ugurbil; 1U54MH091657), funded by the 16 NIH Institutes and Centers that support the NIH Blueprint for Neuroscience Research; and by the McDonnell Center for Systems Neuroscience at Washington University. Glioblastoma imaging and segmentation data were accessed through The Cancer Imaging Archive from the UPENN-GBM collection.

## AI Disclosure

AI-based tools were used to assist with drafting, condensation, and language editing under the authors’ direction; the authors reviewed all content and take full responsibility for it.

